# How do red-eyed treefrog embryos sense motion in predator attacks? Assessing the role of vestibular mechanoreception

**DOI:** 10.1101/634899

**Authors:** Julie Jung, Su J. Kim, Sonia M. Pérez Arias, James G. McDaniel, Karen M. Warkentin

## Abstract

The widespread ability to alter hatching timing in response to environmental cues can serve as a defense against threats to eggs. Arboreal embryos of red-eyed treefrogs, *Agalychnis callidryas*, hatch up to 30% prematurely to escape predation. This escape-hatching response is cued by physical disturbance of eggs during attacks, including vibrations or motion, and thus depends critically on mechanosensory ability. Predator-induced hatching appears later in development than flooding-induced, hypoxia-cued hatching; thus, its onset is not constrained by the development of hatching ability. It may, instead, reflect the development of mechanosensor function. We hypothesize that vestibular mechanoreception mediates escape-hatching in snake attacks, and that the developmental period when hatching-competent embryos fail to flee from snakes reflects a sensory constraint. We assessed the ontogenetic congruence of escape-hatching responses and an indicator of vestibular function, the vestibulo-ocular reflex (VOR), in three ways. First, we measured VOR in two developmental series of embryos 3–7 days old to compare with the published ontogeny of escape success in attacks. Second, during the period of greatest variation in VOR and escape success, we compared hatching responses and VOR across sibships. Finally, in developmental series, we compared the response of individual embryos to a simulated attack cue with their VOR. The onset of VOR and hatching responses were largely concurrent at all three scales. Moreover, latency to hatch in simulated attacks decreased with increasing VOR. These results are consistent with a key role of the vestibular system in the escape-hatching response of *A. callidryas* embryos to attacks.

Red-eyed treefrogs’ hatching responses to predator attacks, vibration playbacks, and egg-jiggling appear when vestibular function develops. Ear development may be a key limiting factor in the onset of mechanosensory-cued hatching.

## INTRODUCTION

Hatching is an essential embryo behavior that mediates the transition between two distinct stages of life, in the egg and post-hatching environments, when developing animals are exposed to different risks and opportunities. Variation in either environment can affect when is the best time to hatch. Environmentally cued hatching allows embryos to respond adaptively to their local environment by altering the timing of their hatching (Sih and Moore, 1993; Warkentin, 1995). Recent syntheses reveal that cued hatching responses are phylogenetically widespread (Warkentin, 2011a). Physical disturbance of eggs is particularly common as a cue for hatching, as observed in invertebrates (Endo et al., 2018; Mukai et al., 2014; Nishide and Tanaka, 2016; Oyarzun and Strathmann, 2011; Whittington and Kearn, 1988), fishes (Martin et al., 2011), amphibians (Buckley et al., 2005; Gomez-Mestre et al., 2008; Goyes Vallejos et al., 2018; Touchon et al., 2011; Warkentin, 1995; Warkentin, 2000; Warkentin, 2011b), and reptiles (Doody, 2011; Doody and Paull, 2013; Doody et al., 2012). Physical disturbance cues can function in antipredator responses, conspecific-cued hatching, host-cued hatching of parasites, and embryo responses to physical conditions (Warkentin, 2011a; Warkentin, 2011b). Physical disturbance may be a particularly useful cue to impending predation of terrestrial eggs, since predators cannot eat eggs without touching and moving them, and terrestrial embryos appear to have less opportunity to receive chemical early warning cues than do aquatic embryos.

To our knowledge, the mechanosensory system mediating hatching responses to physical disturbance cues has not been assessed for any embryos. Indeed, we know relatively little about the developmental onset of mechanoreception, compared to its mature function, across taxa (Hill, 2008). In vertebrates, the predominant motion-detection system is the vestibular system of the inner ear. Otic mechanoreceptors appear during embryonic development in fishes (Becerra and Anadon, 1993; Bever and Fekete, 2002; Haddon and Lewis, 1996), amphibians (Fritzsch, 1996; Quick and Serrano, 2005), chicks (Alsina and Whitfield, 2017; Liang et al., 2010), mice (Fritzsch, 2003; Fritzsch et al., 2002), and humans (Fritzsch et al., 1998). Thus they could potentially mediate mechanosensory-cued hatching. In fishes and amphibians, the lateral line system also develops before hatching (Bever et al., 2003; Hill, 2008; Nieuwkoop and Faber, 1956; Stone, 1933; Thomas et al., 2015) and thus might play a role in mediating cued-hatching responses. Understanding the sensors that mediate cue perception is a key part of understanding any cued behavior and may be particularly crucial early in ontogeny, when both sensory abilities and behavior are changing rapidly. Unlike adults, with fully developed sensory systems, embryos’ ability to respond to particular cue types is constrained by the need for adequate prior development of the relevant sensors. It is essential to identify these sensors and assess their ontogeny in order to determine when developmental changes in behavior reflect the easing of sensory constraints and to understand the information available to embryos at different developmental stages. Understanding sensory ontogeny will also facilitate inquiry into other sources of developmental changes in behavior, such as ontogenetic adaptation of decision rules (Warkentin et al., 2019).

Red-eyed treefrog embryos, *Agalychnis callidryas* (Cope 1862), are tractable study organisms for research on predator-induced, mechanosensory-cued hatching of terrestrial eggs. Females lay gelatinous egg clutches upon leaves and other substrates overhanging ponds, such that hatching tadpoles generally fall into the water below as soon as they hatch (Gomez-Mestre et al., 2008; Pyburn, 1970). Thus hatching, for these embryos, entails a change from arboreal to aquatic habitats with concomitant changes in selection pressures and potential predators (Warkentin, 1995). Predation is the largest cause of mortality for *A. callidryas* embryos monitored at ponds in Costa Rica and Panama, and attacks during the last third of the typical undisturbed embryonic period induce rapid escape-hatching responses of embryos (Gomez-Mestre and Warkentin, 2007; Warkentin, 1995; Warkentin, 2000). Embryos hatch in response to physical disturbance by predators and, at least for snake attacks, playback of recorded attack-vibrations is sufficient to elicit premature hatching (Hughey et al., 2015; Warkentin, 2005). Embryos can hatch within seconds by rapidly releasing hatching enzymes to digest a small hole in the membrane, then squeezing through it (Cohen et al., 2016). Embryos also hatch prematurely in response to flooding, cued by hypoxia (Warkentin, 2002), and drying, based on unknown cues (Salica et al., 2017).

We recently discovered that the developmental onset of hatching responses to hypoxia and mechanosensory cues differs in *A. callidryas* (Warkentin et al., 2017). Specifically, there is a period of development when embryos are competent to hatch, as demonstrated by their consistent hatching response to strong hypoxia and the presence of hatching gland cells, yet still unresponsive to mechanosensory disturbance cues or natural predators (Cohen et al., 2019; Warkentin et al., 2017). Up to 10% of eggs laid can be consumed during this period (Gomez-Mestre and Warkentin, 2007; Warkentin, 1995; Warkentin, 2000; Warkentin et al., 2017), suggesting that an earlier onset of escape-hatching responses to predators could be beneficial. The existence of this hatching-competent but unresponsive-to-predators period indicates that something beyond hatching ability limits the onset of the anti-predator response (Warkentin et al., 2017). The survival cost of early hatching decreases gradually, over days, not hours, of development (Warkentin, 1995; Warkentin, 1999; Willink et al., 2014); this suggests that changes in adaptive embryo decisions are unlikely to impose a narrow developmental limit on the onset of the anti-predator response. Instead, the rapid developmental increase in response to a simulated attack cue, from 0–100% hatching over a few hours (Warkentin et al., 2017), suggests a sensory constraint may limit embryos ability to detect attacks. Thus, as an initial step to assess what sensory system mediates vibration perception in attacks, we looked for ontogenetic congruence of sensor development and the onset of the escape-hatching behavior.

We hypothesize that *A. callidryas* embryos use inner ear mechanoreceptors to sense motion cues. If the vestibular system is the primary mechanism by which red-eyed treefrog embryos sense the physical disturbance of their egg clutches, and is required to perceive predator attacks, its development may limit the onset of escape-hatching responses in this context. To assess this, we looked for developmental correlations between escape hatching behavior and a marker of vestibular function, the vestibulo-ocular reflex (VOR). This reflex generates eye movements that compensate for head movement, producing a more stable image in the retina (Straka, 2010). VOR can be used as a convenient behavioral indicator of vestibular function, as it depends critically on vestibular system development (Jen, 2009). The development of roll-induced VOR has been extensively studied in the frog *Xenopus laevis* (Horn et al., 1986a; Horn et al., 1986b; Horn, 2006; Horn and Gabriel, 2011; Rayer and Horn, 1986), the fish *Oreochromis mossambicus* (Sebastian et al., 2001; Sebastian and Horn, 1999), and the salamander *Pleurodeles waltl* (Gabriel et al., 2012), demonstrating that VOR amplitude is a sensitive indicator of vestibular function. In these aquatic species, modifications of vestibular input either by vestibular lesions (Horn et al., 1986b; Rayer and Horn, 1986; Schaefer and Meyer, 1974) or by altered gravitational conditions during critical periods of vestibular system development (Gabriel et al., 2012; Horn, 2006; Horn and Gabriel, 2011; Sebastian et al., 2001) reliably lead to significant reductions in VOR.

We began by quantifying the basic ontogeny of VOR in *A. callidryas* embryos. We predicted that the developmental onset of VOR (increase in magnitude from absent to consistently strong) would align with the previously documented ontogeny of *A. callidryas’* escape-hatching success in predator attacks. In attacks by egg-predatory snakes and wasps, we have never observed hatching before about 2.5 days prematurely. In our Panamanian study population, escape-hatching success is zero at age 3 d, present but relatively low and variable at 4 d, and consistently high thereafter, with spontaneous hatching in the evening at 6 d (Almanzar and Warkentin, 2018; Hite et al., 2018; Warkentin, 2000; Warkentin et al., 2006a). In Warkentin’s 1990s research on the Osa Peninsula, Costa Rica, with *A. callidryas* developing more slowly under cooler conditions, predator-induced hatching began at age 5 d, with spontaneous hatching in the evening at 7 d (Gomez-Mestre and Warkentin, 2007; Gomez-Mestre et al., 2008; Warkentin, 1995). Accordingly, if a strong VOR were present at the onset of hatching competence at 3 d, vestibular system development would be implausible as a constraint underlying the later onset of attack-induced hatching. Moreover, assuming that VOR is a reliable marker for vestibular function, if VOR onset were clearly later than the onset of escape responses to attacks (e.g., not present until 5 d), it would reject a key role for the vestibular system in sensing attacks.

Next, focusing on the developmental period of greatest variation in VOR and hatching response, we directly compared the hatching responses of egg clutches to vibration playback with the VOR of a subset of hatched and unhatched individuals from each clutch. During this period of high variation, and potentially rapid developmental change, we predicted a positive relationship between magnitude of VOR and hatching response, with some threshold value below which vestibular function is insufficient to cue hatching.

Finally, we applied a simulated predator attack cue to individual eggs to compare hatching responses of embryos to their VOR, as clutches developed. Here as well, we predicted that the onset of vestibular function would match the onset of hatching responses. A lack of correlation of VOR magnitude and hatching response across either clutches or individuals, during the period of high variation and response onsets, would suggest that these are developmentally independent events that simply happen to occur during the same general period. The presence of correlated VOR and hatching responses both among clutches and among individuals would be consistent with a functional linkage.

## METHODS

### Egg clutch collection and care

We collected 0–3 d old *A. callidryas* egg clutches and the leaves on which they were laid from the Experimental Pond in Gamboa, Panama (9.120894 N, 79.704015 W). Clutches were brought to a nearby ambient temperature and humidity laboratory at the Smithsonian Tropical Research Institute, mounted on plastic cards for support, positioned over aged tap water in plastic cups, and misted with rainwater frequently to maintain hydration. Tests of hatching responses were conducted in the ambient-conditions laboratory, and individual hatchlings were tested for VOR in an adjacent air-conditioned laboratory. All embryos used were morphologically normal, in developmental synchrony with siblings in their clutch, and in intact, turgid eggs at the start of the experiment. Most clutches are laid between 10 pm and 2 am, so we assign embryos to daily age-classes and report developmental timing starting from midnight of their oviposition night (Warkentin, 2002; Warkentin et al., 2005). Across the onset of hatching competence, tested individuals were staged based on morphological markers described in Warkentin et al. (2017). Some specimens were preserved for morphological studies (to be presented elsewhere) and all other hatchlings were returned to their pond. This research was conducted under permits from the Panamanian Environmental Ministry (SC/A-15-14, SE/A-46-15) and approved by the Institutional Animal Care and Use Committees of Boston University (14-008) and the Smithsonian Tropical Research Institute (2014-0601-2017).

### Measurement of the vestibulo-ocular reflex (VOR)

We measured roll-induced VOR of newly hatched tadpoles or manually decapsulated embryos (henceforth, collectively ‘hatchlings’) using a custom-built, Arduino-based, portable tadpole rotator (Fig. S1, J. G. McDaniel, Adrian Tanner, and K. M. Warkentin; Boston University Engineering Products Innovation Center). The rotator smoothly turns a shaft at the push of a ‘clockwise’ or ‘counterclockwise’ button and was programmed for 15° rotational increments. A conditioning mass and rubber plate mounted on the shaft limit vibration transfer from the motor to the test animal, and a printed plastic cup glued to the rubber plate enables field replacement of the animal interface. To hold hatchlings, we mounted a section of plastic pipette in the center of the cup, in line with the rotator shaft, using silicone seal. The hatchling chamber was 13.5 mm long and 3 mm in internal diameter, with a slight widening at the mouth so as not to restrict eye motion, and horizontally leveled in relation to gravity. To test a hatchling, the chamber was filled with aged tap water and the animal was backed into it using a transfer pipette or length of tubing on a syringe, positioning its snout just within the tube. No anesthesia was necessary and individuals could be tested within minutes of hatching.

The chamber was surrounded by a light diffuser and illuminated on both sides by LED lights (Panasonic 9W, 100-127V, 90mA), providing a uniform white visual field. It faced a horizontally leveled MPE-65 mm macro lens on a digital camera (Canon D70) with cable shutter release, mounted on a focusing rail on a tripod. Following Horn (Horn et al., 1986b; Horn and Sebastian, 1996), we rolled hatchlings about their body axis 180° in each direction, photographing them in frontal view each 15° (Movie S1). We continuously observed hatchlings on the camera view-screen, manually applied rotation increments, and took each photograph as soon as body and eye rotation had stopped, to minimize testing time. Most animals remained immobile through each 180° roll sequence; for those that moved more than their eyes, we restarted the sequence from 0° to obtain a continuous series of measurements. From each photograph, we measured right and left eye angle and body axis angle using ImageJ (Schneider et al., 2012). From each angular measurement series, we constructed an individual VOR curve using a sine-fitting function in Python (Version 2.7.9, Build 1, Python Software Foundation). We assessed the curve fit and calculated the VOR amplitude from the sine function. The peak-to-peak amplitude of the curve corresponds to the hatchling’s VOR magnitude (Fig. 1).

**Fig. 1.**
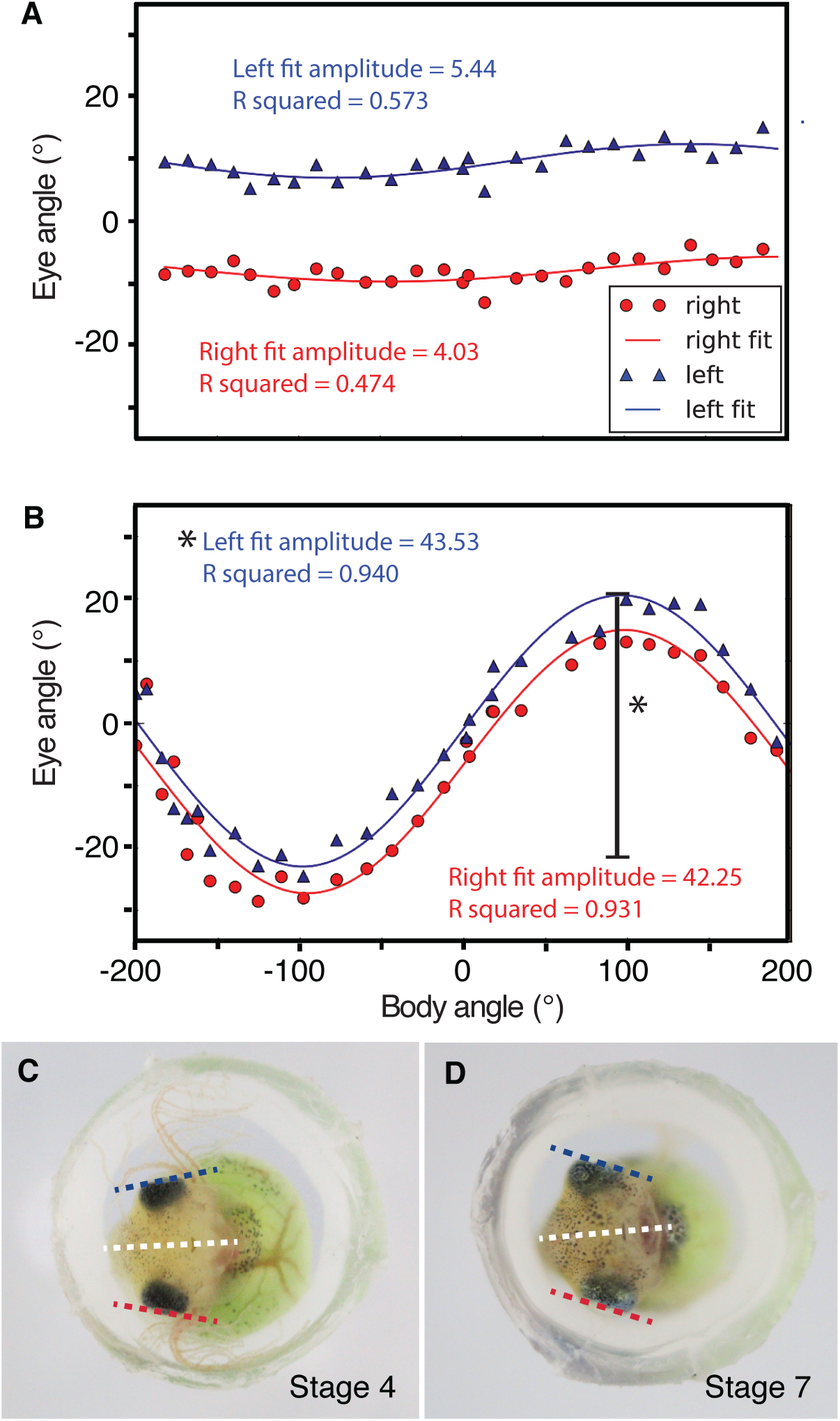
Example vestibulo-ocular reflex (VOR) curves for *Agalychnis callidryas* hatchlings. (A, B) Data points show eye angle (left in blue, right in red) plotted against body angle and lines are best fit curves for each eye. The peak-to-peak amplitude (*) of the curve measures the magnitude of VOR. (C, D) Frontal view of hatchlings tested, showing left (blue) and right (red) eye angles relative to the body axis (white). (A) Low VOR for a hatchling of age 3.6 d (C). (B) Clear VOR in a hatchling of age 4.5 d (D).

We visually checked each sine curve fit and rejected those that did not meet the following criteria: 1) curve fits of the two eyes show similar wavelengths, are horizontally aligned, and have parallel or near-parallel waveforms, 2) the wavelength is plausible for VOR, with zero crossing at or near the zero body angle, and 3) eye rotation is opposite to body rotation (i.e., curve is not upside-down). Individuals whose curve fits failed one or more of these VOR criteria (Fig. S2) were considered to have a VOR of zero. Of 406 hatchlings tested, 92 failed the VOR curve fit criteria (N = 4 of 36 in series **Ia** below, N = 38 of 89 in **Ib**, N = 19 of 169 in **II**, N = 31 of 112 in **III**).

#### I. Ontogeny of vestibular function

First, to determine the basic ontogenetic timing of the onset of vestibular sensory function in *A. callidryas*, we measured the VOR of embryos at different ages (**Ia)**. From 19–25 June 2014, we tested VOR daily across the plastic hatching period, in the afternoon of each day (13:27– 17:09 h), using a set of non-sibling embryos at each age (N = 7, 10, 10, and 9 hatchlings, at ages 3–6 d respectively; total N = 36 hatchlings from 14 clutches). Second, to assess how VOR varied among and within egg clutches across development, we tested developmental series of five clutches, from 10–20 August 2014 (**Ib)**. We concentrated our sampling in the period of greatest change, testing ∼ three siblings per age at 6 h intervals from 3.75–4.75 d, with a final sample at 5.75 d (total N = 88 hatchlings; N = 15, 16, 15, 15, 15, 12 individuals per age group).

We removed each individual egg from its clutch just prior to VOR testing, placed it in a small dish, and gently rolled and jiggled it with a blunt probe to induce hatching. The youngest embryos were unresponsive to this stimulus and, instead, manually decapsulated with fine forceps under a dissecting microscope. Hatchlings were tested for VOR within 3 minutes of leaving their egg capsule.

#### II. VOR and hatching response in vibration playback to whole clutches

To assess if variation in VOR and the hatching response are related, across the period of high variation in both traits, we paired vibration playbacks to 36 clutches with VOR measurements on a subset of embryos from each clutch, from 26 June to 21 July 2015. We tested clutches at ages 3.7–4.9 d and stages 1–7 (Warkentin et al., 2017), from before any hatching response to vibration until responses became fairly strong. To focus on fine-scale developmental changes and avoid age-imprecision due to variation in oviposition timing, we report results based on stage. Compared to 2014, development tended to be accelerated under the warm El Niño conditions in 2015 (Warkentin et al., 2017).

We played a synthetic low-frequency vibration stimulus (Fig. 2A) designed to elicit very high hatching, based on prior playbacks to 5-d-old clutches (Caldwell et al., 2009; Warkentin et al., 2006b). We generated noise in Matlab and filtered it using a custom script, playback.m (available upon request), to compensate for nonlinearities in the shaker transfer function and generate a frequency distribution resembling that of snake attacks (Caldwell et al., 2009), with high energy below 60 Hz and intensity dropping off above that (Fig. 2C). To test our match to the desired frequency distribution, we recorded playbacks of the stimulus embedding a small (0.14 g) AP19 accelerometer (AP Technology International B.V., Oosterhout, The Netherlands) within a clutch. Accelerometers added ∼5% to the mass of each clutch, such that test clutches remained within the natural range of interclutch mass variation (Warkentin 2005). Transduced vibrational signals were powered/amplified by an APC7 signal conditioner and digitized with an external sound card (MSE-U33HB; Onkyo USA, Saddle River, NJ, USA). The output was recorded using Raven Pro 1.3 bioacoustics software (Cornell University Laboratory of Ornithology, Ithaca, NY, USA) on a Macbook Pro computer. The intensity of frequencies below 20 Hz was limited by shaker capabilities. The base temporal pattern consisted of 0.5 s pulses of vibration, with roughly rectangular amplitude envelopes, separated by 1.5 s intervals of silence (Fig. 2B). This was divided into pulse-groups consisting of 10 pulses separated by 30-s gaps of silence (Fig. 2B). We included a three-pulse “primer” plus 30-s gap before the repeating10-pulse pattern began, since this element also increases hatching response (Fig. 2B, Jung, Guo, McDaniel and Warkentin, unpublished data).

**Fig. 2.**
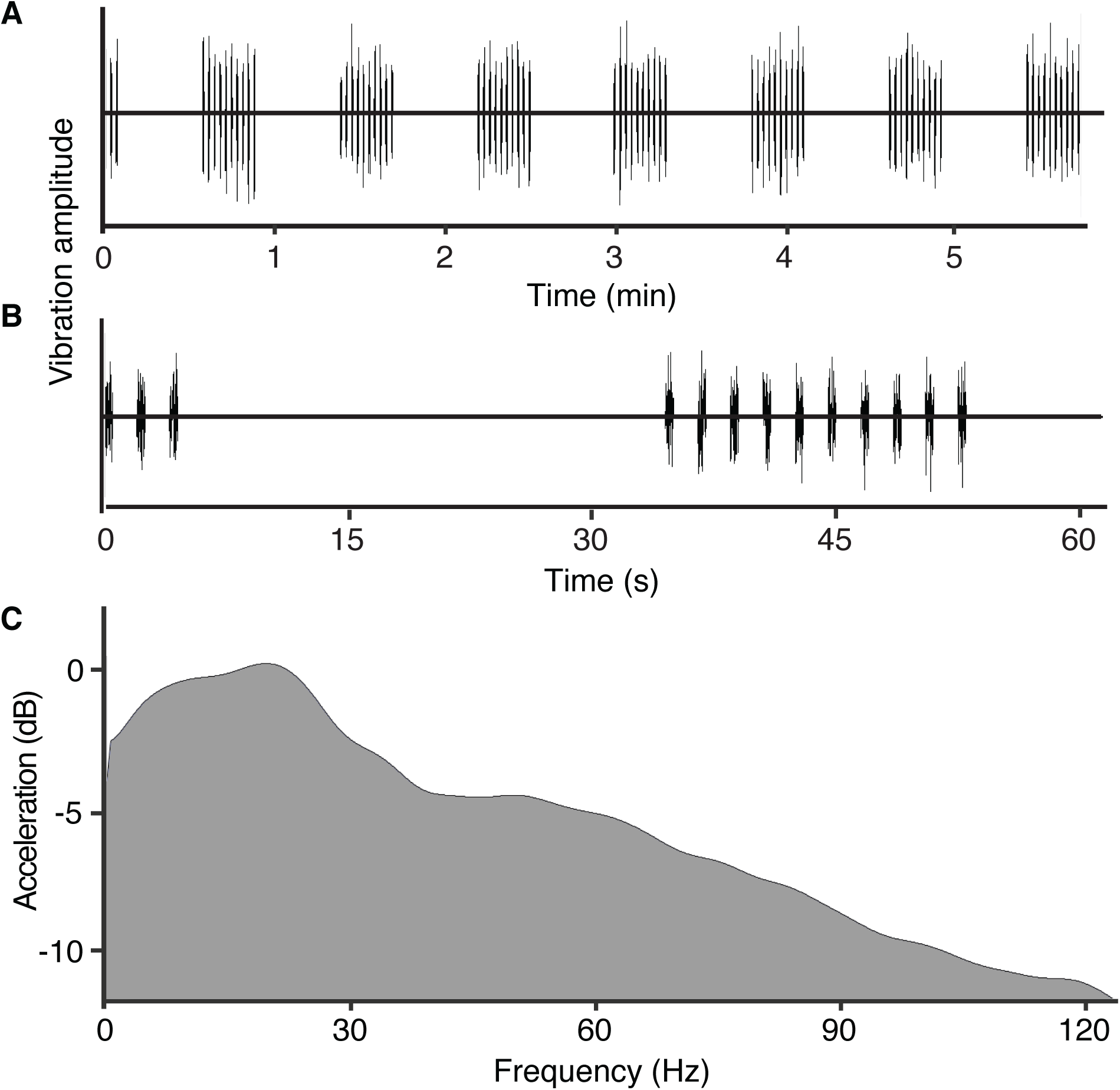
Vibration playback stimulus. (A) The entire stimulus included 7 pulse groups divided by 30 s gaps plus a 3-pulse primer before the first 30 s gap. (B) The first minute of the stimulus, including the primer and one pulse group, comprised of ten 0.5 s pulses of vibration, separated by 1.5 s intervals. (C) Frequency spectrum of acceleration, normalized from peak power. This stimulus induces near 100% hatching in 5-day-old clutches.

Playback methods followed Caldwell et al. (Caldwell et al., 2009; Caldwell et al., 2010). Stimuli (Fig. 2A) were presented through an array of blunt metal tines inserted among eggs (Fig. S3, Movie S2) attached via a rigid post to an electrodynamic minishaker (Model 4810; Brüel & Kjær, Nærum, Denmark). Shaker output was controlled by Audacity 2.1.0 (Free Software Foundation, Boston, MA) on a 2014 MacbookAir, via a custom-made amplifier designed to have a flat frequency response from DC to 5 kHz (E. Hazen, Boston University Electronic Design Facility). Playback clutches on their plastic cards were mounted on a flat-sided plastic stand (∼1.5 kg), then carefully slid forward so the tines entered the clutch between eggs. We used only healthy clutches that fit within the tine field, and tines were rinsed with rainwater between trials. We watched for any hatching induced by the set-up procedure (only 3 individuals, from 3 clutches), then allowed five hatching-free minutes for acclimation before starting the playback. For playback, the shaker moved the tines up and down, so eggs were shaken vertically, and hatched tadpoles fell into a tray of water below the clutch.

For each trial, we counted the embryos that hatched during the playback period and 5-min of post-playback observation. We then immediately (within 5–10 minutes) measured VOR of a subset of 3 hatchlings per clutch that had hatched in response to playback, unless fewer had hatched. To check for hatching competence of the remaining eggs, after post-playback observation, we manually stimulated eggs, rubbing and jiggling them with a blunt metal probe, for about two minutes, then submerged any unhatched eggs in hypoxic water. Any embryos that failed to hatch under manual stimulation and hypoxia were considered not competent to hatch, and excluded from the count of test individuals in calculations of proportion hatched per clutch (proportion excluded = 0.078 ± 0.022, mean ± s.e.m. across clutches). We measured VOR of 3 additional hatchlings that hatched in response to either manual stimulation or hypoxia, but not vibration playback; numbers of manually-stimulated and hypoxia-cued hatchings tested for VOR varied among clutches (total of 3, unless fewer remained after playback). We staged all VOR-tested hatchlings (N = 143 hatchlings total) from their frontal photos following a staging system adapted from Warkentin et al. (2017) (Fig. 3).

**Fig. 3.**
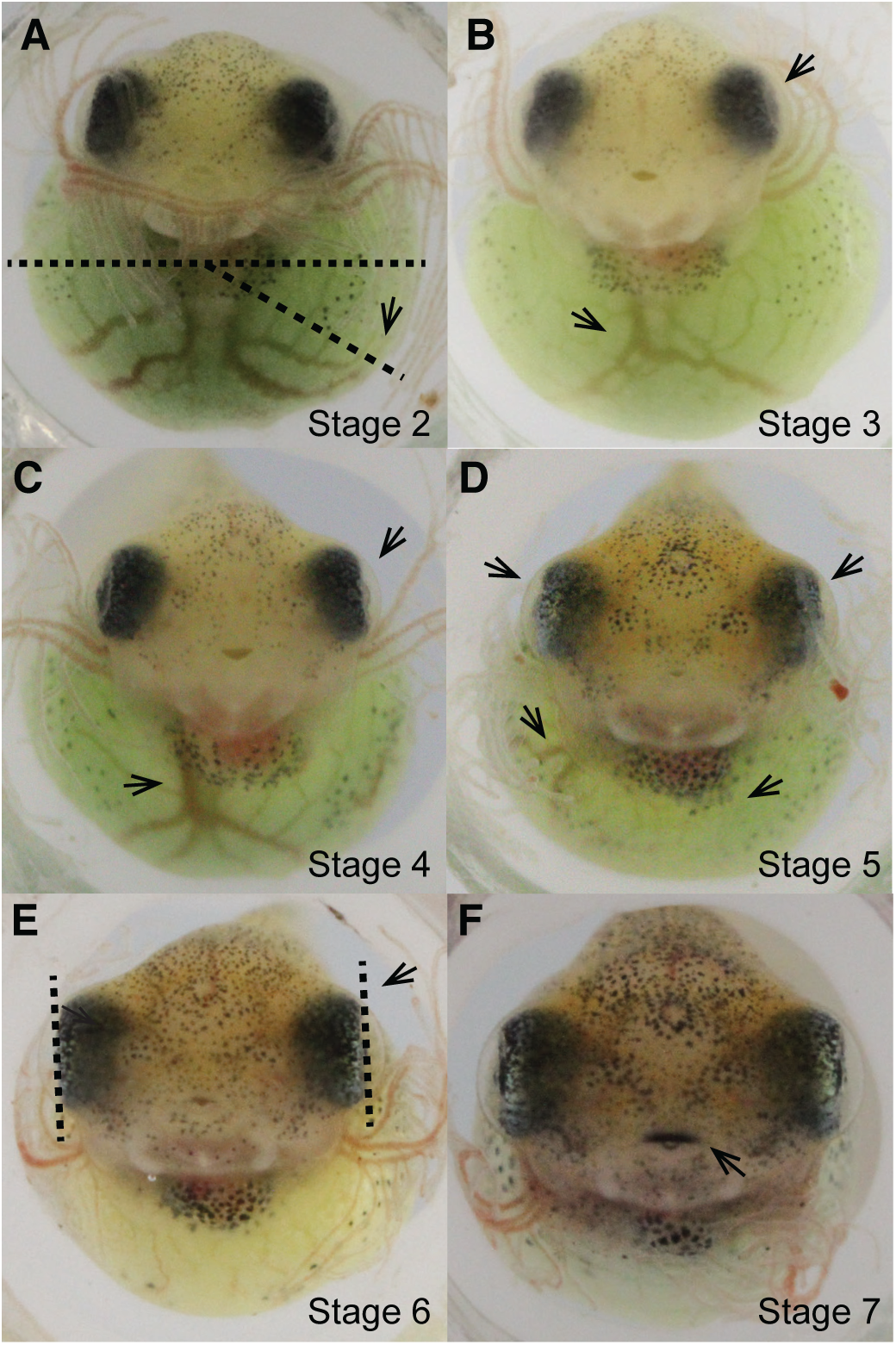
Stages in the development of *Agalychnis callidryas* embryos through the onset of hatching. Staging traits are adapted from Warkentin et al. (2017) and all visible in frontal view. (A) Stage 2 – Melanophores extend at least halfway down sides over yolk. Two dominant veins on yolk surface enter heart separately, fairly symmetrically under center of heart. (B) Stage 3 – Two dominant veins on yolk surface join to enter heart as a single vessel. Cornea close to lens and granular or slightly cloudy, partially obscures view of lens. (C) Stage 4 – Cornea clear and well separated from lens; lens readily visible. Yolk vein enters medially below heart, within edge of cement gland in direct frontal view, veins present in yolk posterior to heart. (D) Stage 5 – Yolk vein enters heart dextrally, at or lateral to cement gland in direct frontal view, yolk posterior to heart clear of veins. Eyes angled in dorsally. (E) Stage 6 – Eyes parallel in frontal view, not angled in dorsally. Beaks unkeratinized. (F) Stage 7 – Edge of upper and/or lower beaks keratinized, head narrower than yolk.

#### III. VOR and hatching response to simulated attack on individual embryos

To examine the correlation between hatching responses to physical disturbance cues and vestibular function on an individual level, we assessed both traits in developmental series of embryos across the onset of mechanosensory-cued hatching. To assess hatching responses of embryos to a simulated attack, we removed individual eggs from their clutch, placed each in a petri dish with a drop of water, and manually jiggled them with a moistened blunt metal probe, alternating 15 s of stimulation and 15 s of rest for 5 min or until the egg hatched (Warkentin et al., 2017) (Movie S3). We tested two embryos per clutch from 11 clutches every 3 hours, on August 11–13, 2015. As with vibration playbacks, we observed embryos for 5 min before, during, and after stimulation (15 minutes total), and considered any hatching during and after stimulation (10 minutes) to be a response to the stimulus. All sibships were initially tested for their hatching response to hypoxia and, in most cases, we began testing responses to the egg-jiggling stimulus only after siblings had demonstrated an ability to hatch; the data on developmental timing of onset of the response to each cue are reported elsewhere (Warkentin et al., 2017). We continued testing each clutch every 3 h until both test embryos had hatched at two time points, thus capturing a range of developmental ages (3.25–4.625 d) and stages (2–7) from those unresponsive to the jiggling cue, through the onset of response, to strongly responsive (total N=112 individuals, 6–18 per clutch). For each hatchling we recorded latency to hatch, from stimulus onset, or failure to hatch after 5 min of post-stimulus observation. We manually decapsulated unhatched embryos, and photographed all animals in frontal view to assess development (from stages 2–8) following a staging system adapted from Warkentin et al. (2017) (Fig. 3). We then immediately measured their roll-induced VOR.

### Statistics

When data met parametric assumptions, we used ANOVAs and Tukey post-hoc tests to find effects and comparisons. Otherwise, we used the Wilcoxon Rank Sum and Wilcoxon Each Pair methods for non-parametric tests of effects and comparisons. In the first developmental assay that examined the hatching response of multiple siblings per clutch (**Ib**), we fit a 4-parameter logistic model, grouped by clutch, and performed an analysis of means for inflection point estimates. In the following assays where we considered multiple siblings per clutch, we analyzed our results using mixed models with clutch as a random effect. To analyze predictors of hatching, we used binomial GLMMs with clutch as a random effect and performed likelihood ratio tests to compare nested models. All statistical tests were carried out in JMP Pro 13 (version 13.2.0, SAS Institute Inc. 2016) or the R statistical environment (version 3.3.3, R Development Core Team 2014, http://www.r-project.org) in RStudio (version 1.1.383, RStudio Team 2015).

## RESULTS

### I. Ontogeny of vestibular function

Across embryos tested at daily intervals, VOR amplitude increased with age (Wilcoxon Rank Sum: χ^2^=16.2797, df=3, P=0.0010, Fig. 4A). VOR did not change significantly from age 4–6 d (Wilcoxon Each Pair, all P>0.4274) but hatchlings tested at age 3 d showed lower VOR than those aged 4–6 d (Wilcoxon Each Pair, all P<0.0014, Fig. 4A). In the second developmental series, with replication within clutches, VOR increased with age in a sigmoidal fashion (R^2^=0.91, Fig. 4B), and clutches varied in inflection point estimates (Analysis of means; upper limit exceeded in clutch 101 and 102, P<0.01; Fig. 4B).

**Fig. 4.**
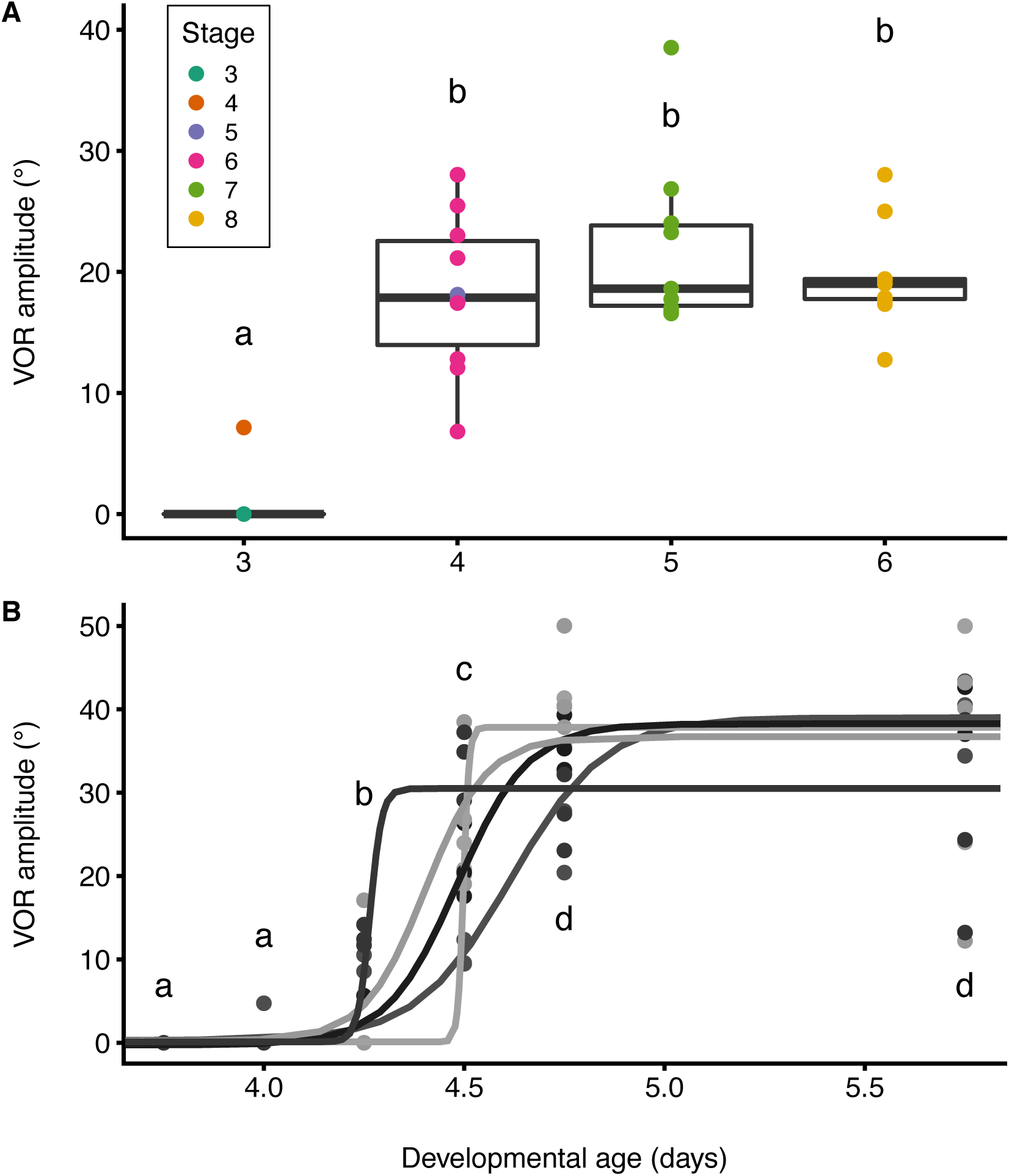
Ontogeny of the vestibulo-ocular reflex (VOR) across embryonic development of *Agalychnis callidryas*. (A) Hatchings were tested at 24 h intervals from age 3–6 d (at approximately 3 pm), using 7–10 non-sibling individuals per age (36 hatchlings tested from 14 different clutches). Individual hatchlings were tested for VOR immediately after hatching or decapsulation and are color-coded by developmental stage. Box plots show medians, interquartile range (IQR), and extent of data to ± 1.5×IQR. Different letters indicate significant differences in VOR amplitudes between ages. (B) Individual hatchlings were tested from developmental series of five clutches (shown in different shades of gray), with ∼3 siblings per test point (total N=88 hatchlings; N=15, 16, 15, 15, 15, 12 per age). Lines show 4-parameter logistic curve fit for each clutch.

### II. VOR and hatching response in vibration playback to whole clutches

Based on post-playback hypoxia testing, all individuals included in VOR analyses (N=169) were able to hatch, but only 63 of them hatched in response to vibration playbacks. VOR amplitude increased significantly across developmental stages (one-way ANOVA, f5,162=79.2953, P<0.0001). Across the first four stages we tested (stages 2–5, Warkentin et al. 2017), no embryos in any clutches hatched in response to vibration playbacks and VOR was consistently low (5.3±1.0°, mean±SE, here and throughout; N=22 hatchlings, 8 clutches; Fig. 5A). Compared to VOR at stages 2–5, VOR was higher at stage 6 (N=63 hatchlings, Tukey test from one-way ANOVA, P<0.0001) and stage 7 (N=84 hatchlings, P<0.0001, Figure 5A). Up until stage 5, no individuals hatched. At stage 6, vibration-cued hatching began, but the low hatching response rates within clutches were not significantly higher than zero at earlier stages (Tukey test from one-way ANOVA, P>0.3809); clutch hatching rates at stage 7 were significantly higher than at all prior stages (P<0.0001).

**Fig. 5.**
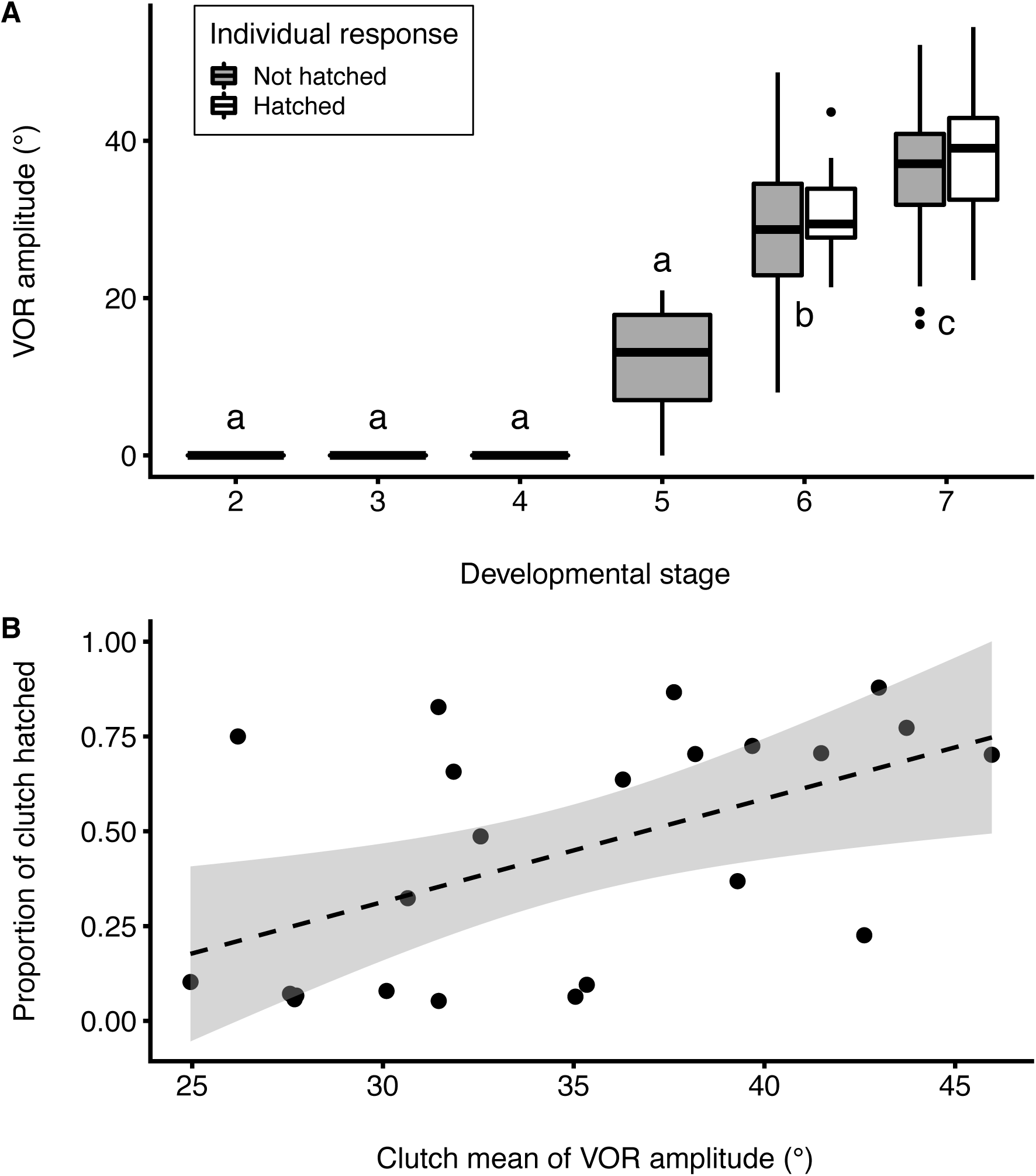
Relationships among vestibulo-ocular reflex (VOR) amplitude, development, and hatching response of *Agalychnis callidryas* embryos to vibration playbacks. (A) Filled and unfilled box plots indicate VOR of individuals (N=169) that did not hatch or hatched, respectively, in response to vibration playbacks to whole clutches across development. Box plots show medians, interquartile range (IQR) and extent of data to ± 1.5×IQR, and outliers as points. Different letters indicate significant differences between stages. (B) Total proportion hatched, for the 23 clutches from which at least one embryo hatched, plotted against clutch mean VOR amplitude measured from 169 hatchlings (4–6 individuals per clutch). Dashed line indicates the linear regression fit, shading indicates 95% confidence interval.

Considering all tested individuals, those that hatched in playback had a significantly higher VOR than individuals that did not hatch in playback, but hatched in response to manual stimulation or hypoxia (Mixed Model, VOR Amplitude ∼ Hatching with Clutch as a random effect: χ^2^=4.8028, P=0.02841). For the subset of 23 clutches where hatching occurred, individuals that hatched in playbacks tended to have higher VOR than siblings that did not hatch, but the difference was not significant (Mixed Model, χ^2^=3.0898, P=0.07878). However, individuals that hatched only in response to hypoxia had significantly lower VORs than those that hatched in response to playbacks (Mixed Model, χ^2^=6.2563, P=0.0438). For clutches with hatching, we compared mean VOR of each clutch (from 5.61±0.14 individuals per clutch, range 4–6) with proportion hatched. Proportion hatched per clutch increased with mean VOR (Figure 5B, linear regression, F1,21=8.0252, P=0.0100); no individuals hatched with VOR less than 21.38° and mean VOR of those that hatched in response to playback was 35.92°±0.95°.

### III. VOR and hatching response to simulated attack on individual embryos

For individual embryos (N=112) subjected to a simulated attack, VOR increased in magnitude across embryonic developmental stages (Mixed Model, χ^2^=96.215, P<2.2e-16, Fig. 6A) and varied among clutches (Mixed Model, χ^2^=8.3355, P=0.003888). No stage 2 embryos hatched. Hatching in response to individual egg-jiggling began at stage 3 with a hatching rate of 20.8% (Fig. 6B), which is when some embryos started showing a measurable VOR (Fig. 6A). By stage 4, almost half the embryos hatched (47.4%) and by stage 7, all embryos tested hatched in response to the jiggling cue (Fig. 6B).

**Fig. 6.**
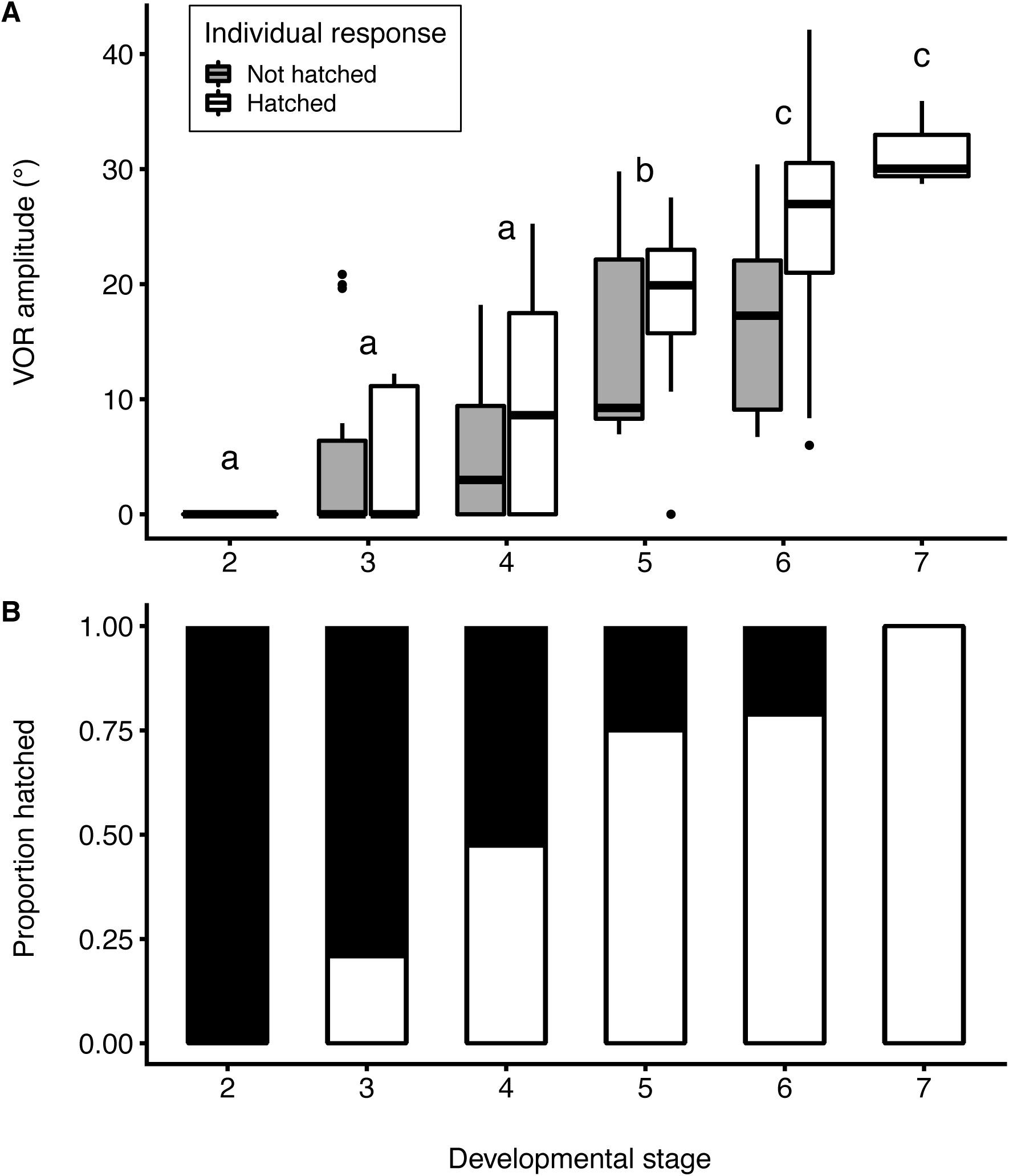
Ontogeny of vestibulo-ocular reflex (VOR) and hatching responses in *Agalychnis callidryas*. (A) Ontogeny of VOR across stages, from developmental series of 11 egg clutches, with two siblings tested per time point (N = 112 individuals). Different letters indicate significant differences between stages. Box plots show medians, interquartile range (IQR) and extent of data to ± 1.5×IQR, and points show outliers. Filled and unfilled box plots indicate embryos that did not hatch and hatched, respectively, in response to manual egg jiggling. (B) Ontogeny of hatching response to manual jiggling of individual eggs. Proportion hatched (white boxes) is of individual embryos tested per stage.

Both developmental stage and VOR amplitude were significant and strong predictors of hatching (Fig. 7A-B, binomial GLMM), and the model incorporating both variables was better than models with either one alone (AIC values 120 vs. 122 and 125). More developed embryos with greater VOR were more likely to hatch, with hatching response increasing 14% for every stage (χ^2^=13.285, P=0.02085) and 24% for every 10 degrees of VOR amplitude (χ^2^=11.951, P=0.0005461, Fig. 7A-B). However, the 61 embryos that hatched in response to egg-jiggling included 7 individuals with no detectable VOR, ranging from stage 3 to 5 (Fig. 8).

**Fig. 7.**
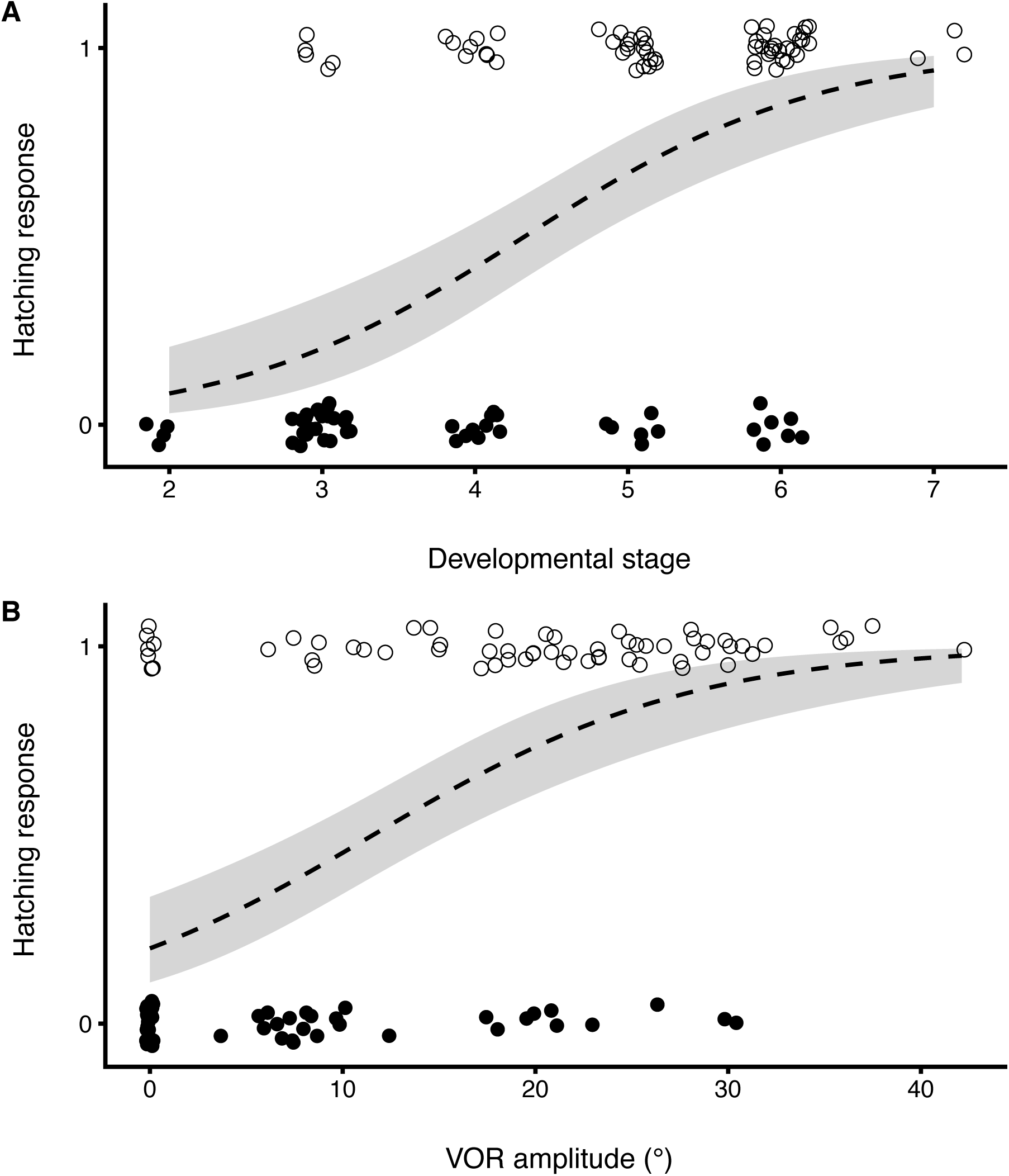
Effect of (A) developmental stage and (B) vestibulo-ocular reflex (VOR) amplitude on hatching response of *Agalychnis callidryas* embryos to egg jiggling. Values of 0 (unhatched, filled circles) and 1 (hatched, open circles) are jittered vertically to show data points. Integer values (developmental stages) are jittered, while continuous individual measurements (VOR amplitude) are not. Dashed lines are predicted fits from binomial generalized linear mixed models; shading indicates the 95% confidence interval.

**Fig. 8.**
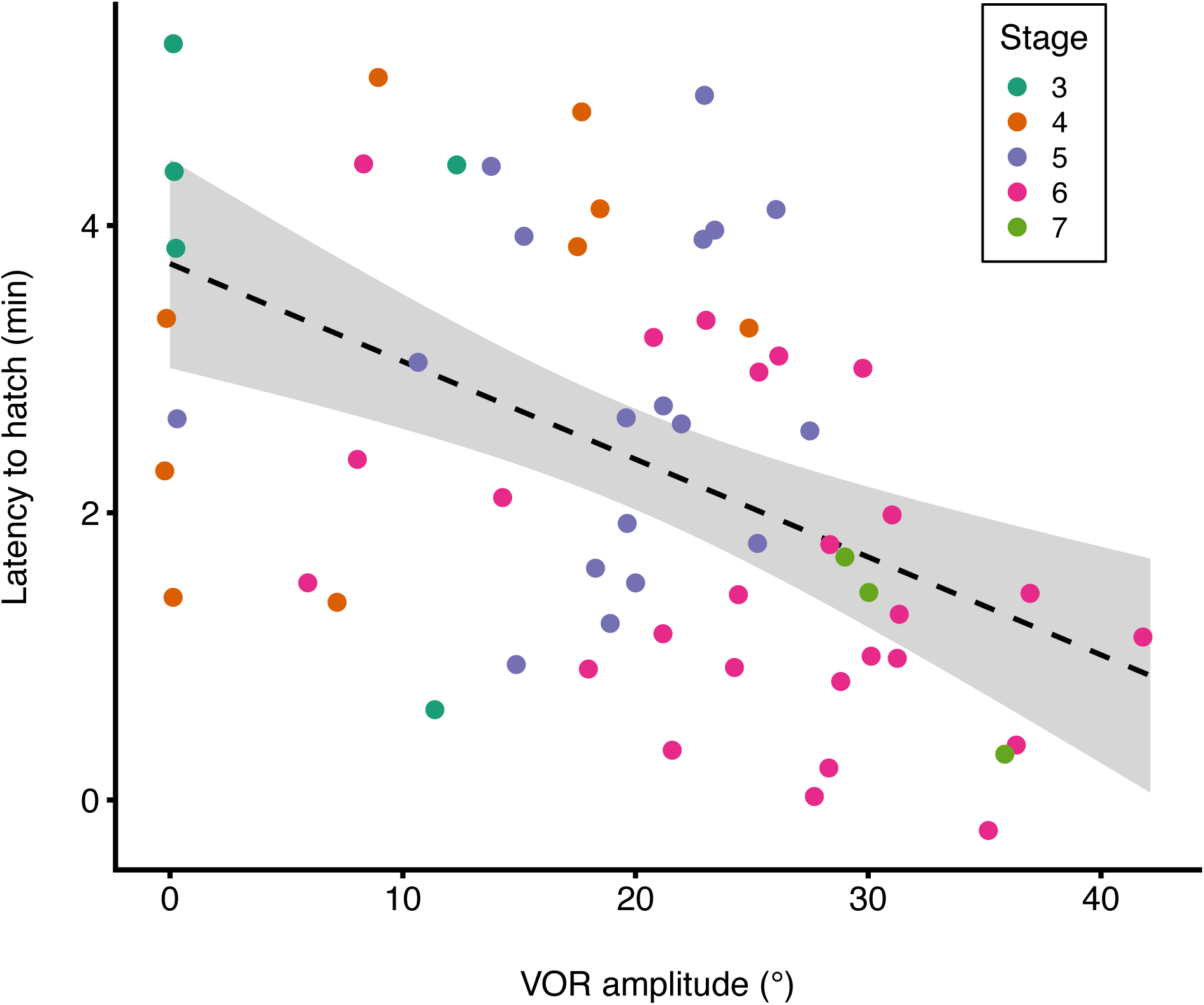
Latency of *Agalychnis callidryas* embryos to hatch in response to egg jiggling, in relation to vestibulo-ocular reflex (VOR) amplitude. Data are for N=61 embryos tested individually. Developmental stages are indicated by color. Dashed line indicates linear regression fit; shading shows the 95% confidence interval.

Considering the subset of animals that hatched in response to egg jiggling, their latency to hatch decreased with VOR amplitude (Fig. 8, Latency ∼ VOR with Clutch as a random effect, χ^2^=16.55, P=4.738e-5). If we add stage and the interaction between stage and VOR into the model, there is a main effect of stage (χ^2^=13.3925, P=0.009509), and an interaction effect (χ^2^=12.0126, P=0.017258), but no main effect of VOR (χ^2^=1.3143, P=0.251613). Closer examination of the interaction indicates a significant VOR effect only at stage 6 (χ^2^=5.0734, P=0.0243), but note that sample sizes were lower at other stages (in order from stage 3, N=5, 9, 18, 26, 3).

## DISCUSSION

Embryos use physical disturbance (egg motion) as a cue to hatch among fishes (Martin et al., 2011), amphibians (Buckley et al., 2005; Gomez-Mestre et al., 2008; Goyes Vallejos et al., 2018; Touchon et al., 2011; Warkentin, 1995; Warkentin, 2000; Warkentin, 2011b), and reptiles (Doody, 2011; Doody and Paull, 2013; Doody et al., 2012), as well as many invertebrates (Endo et al., 2018; Mukai et al., 2014; Oyarzun and Strathmann, 2011; Tanaka et al., 2016; Whittington and Kearn, 1988). However, the specific sensors mediating the environmentally cued hatching responses of embryos are entirely unknown. We examined the role of the vestibular system – the general vertebrate motion sensor – in the escape-hatching response of red-eyed treefrogs. In four experiments, at population, clutch, and individual levels, we found developmental congruence between the onset of the VOR and the escape-hatching response to real and simulated predator attack and vibration playbacks, consistent with our hypothesis that the vestibular system plays a key role in mediating mechanosensory-cued hatching.

### VOR as an indicator of vestibular system function

Our tests for ontogenetic congruence of vestibular system function and escape-hatching behavior are based on the vestibulo-ocular reflex (VOR), or eye movements induced by roll and tilt of the body (Horn et al., 2013), which we could measure within minutes of hatching using a tadpole-in-tube rotation protocol. Since input from the vestibular system controls the muscles responsible for VOR, it is well-established that the VOR is not expressed without vestibular system function (Cohen, 1974; Precht, 1976). Moreover, the onset of VOR appears not to be limited by eye muscle development. Extraocular motoneurons develop and establish axonal connections with target eye muscles very early in embryogenesis (Gilland and Baker, 2005; Glover, 2003). In 96 of 406 hatchlings tested, we observed non-VOR-related eye movements (criteria listed in methods) with a measurable magnitude greater than that of individuals with a small but clear VOR (Fig. S2). This indicates that hatchlings, prior to developing a working VOR, can change their eye angle – just in a way that does not match up with their body rotation. The data from these individuals supports that the onset of VOR is not limited by when embryos become physically capable of moving their eyes. Moreover, the presence of non-VOR-related eye movements motivate our criterion rejecting individuals with non-parallel curve fits. The eye muscles that enable the VOR receive their information from both vestibular organs (Precht, 1976). In *Xenopus*, complete unilateral vestibular lesions and selective lesions of each utricular organ reduce the VOR of both eyes (Horn et al., 1986b). Thus, we considered non-parallel curves for the two eyes to indicate non-VOR-related eye movements (Fig. S2A).

### Ontogenetic congruence of VOR and mechanosensory-cued hatching

When we began this work, we knew that hatching ability does not limit the onset of hatching responses to predator cues, because younger embryos demonstrate hatching competence in response to strong hypoxia (Cohen et al., 2019; Warkentin et al., 2017). Moreover, the rapid developmental increase in hatching response to egg-jiggling – contrasted with the much slower developmental decrease in the costs of early hatching – suggests that some sensory constraint imposes a developmental limit on the onset of the anti-predator response (Warkentin et al., 2017). We performed four experiments to examine the role of vestibular mechanoreception in embryos’ risk assessment by comparing the ontogeny of responses at a population level, at a clutch level, and at an individual level.

First, at a population level, we found that the developmental onset of the vestibulo-ocular reflex in red-eyed treefrog embryos (individually in series **Ia** and across clutches in series **Ib**) is congruent with the documented onset of escape-hatching responses to predator attacks in the Gamboa population of *A. callidryas* (Almanzar and Warkentin, 2018; Hite et al., 2018; Warkentin, 2000; Warkentin et al., 2006a). If the onset of VOR were clearly before or after the developmental period when predator-induced hatching begins in this population of *A. callidryas*, it would have rejected the hypothesized key role of vestibular system development in enabling the antipredator response. Moreover, clutches appeared to vary slightly in their onset of VOR (**Ib**), congruent with the greater variation in escape-hatching success of clutches attacked when the response first appears, and decreased variation later in development (Gomez-Mestre et al., 2008; Warkentin et al., 2006a).

Next, we examined the ontogeny of VOR in more detail through the period of greatest change (series **II**) and tested its relationship to the hatching response using vibration playbacks to entire clutches. Across the onset of ear function, embryos below a VOR threshold of 21° did not hatch during vibration playbacks, even though they could hatch if flooded. Moreover, clutch hatching response increased with clutch mean VOR at supra-threshold levels. These data are also consistent with a key role of vestibular mechanosensing in mediating vibration-cued hatching.

In our last series (**III**), we compared VOR and the hatching response to simulated attacks on individual embryos, rather than whole clutches, and saw that they were still highly correlated. In addition, embryos with greater VOR hatched more rapidly in response to egg jiggling. Hatching occurred developmentally earlier in response to targeted jiggling cues (**III**) than in response to whole-clutch vibration playback (**II**), at stage 3 vs. stage 6. Moreover, embryos started showing a measurable VOR at earlier developmental stages in the egg jiggling series (**III**), relative to the clutch vibration series (**II**) (compare Fig. 5A vs. 6A). VOR development was correlated with stage, but not perfectly. For instance, some stage 3 animals showed VOR but most did not, and one stage 5 animal lacked VOR, but most showed it. VOR amplitude predicted hatching more strongly than did developmental stage although, controlling for VOR, stage explains some additional variation and vice-versa. This individual-level correlation between vestibular function and hatching is consistent with a role of the vestibular system in mechanosensory-cued hatching.

Across successively finer levels of developmental precision, our results reveal a substantial increase in mechanosensory-cued hatching responses with the development of vestibular function, consistent with a role for this sensory system in mediating the response. In general, the timing of onset of vestibular function is consistent with the onset of escape success in predator attacks (Almanzar and Warkentin, 2018; Hite et al., 2018; Warkentin, 2000; Warkentin et al., 2006a). In our vibration playbacks to clutches, no embryos lacking VOR hatched. In our egg-jiggling developmental series, we found a strong correlation of VOR with increased hatching response and decreased hatching latency. However, some evidence suggests that additional mechanoreceptor systems can also play a role in escape-hatching (Fig. 7B).

### Mechanosensory-cued hatching before vestibular function

Of the 61 embryos that hatched in response to our individual egg-jiggling cue, seven individuals (11%) had no detectable VOR; they hatched an average of 4.85 h before their siblings showed VOR. Hatching of embryos lacking VOR in response to jiggling cues is relatively rare and does not reject a key role of the vestibular system in risk assessment by embryos, given the strength of the relationship between VOR and hatching. However, the occurrence of any mechanosensory-cued hatching prior to vestibular function indicates that vestibular mechanoreceptors are not the only sensors that can mediate hatching when eggs are physically disturbed, at least under some types of disturbance. *A. callidryas* embryos clearly use cues in multiple sensory modalities, including hypoxia (Rogge and Warkentin, 2008) and light level (Güell and Warkentin, 2018), to inform hatching. These embryos might also use multiple mechanosensors, either to perceive different cue components available in attacks and egg-jiggling or as potentially redundant or synergistic sensors of the same cue element. Two other candidate sensor types––lateral line neuromasts and cutaneous mechanoreceptors––may also be relevant to mechanosensory-cued hatching in the egg-jiggling context.

### Other mechanosensory systems

The lateral line is a system of mechanoreceptors that detect movement, pressure gradients, and vibration in fishes and aquatic amphibians (Mogdans and Bleckmann, 2012). The effective stimulus to lateral line is low frequency particle motion of the surrounding fluid, relative to neuromasts distributed on the animal’s surface (Strelioff and Honrubia, 1978; Weeg and Bass, 2002). *A. callidryas* embryos develop a lateral line system on their head, body, and tail by 3 d, well before mechanosensory-cued hatching begins at 4 d (Cohen et al., 2019; Warkentin et al., 2017). However, the number of superficial neuromasts, visualized with the fluorescent vital dye DiAsp (Sigma D-3418), continues to increase through the onset of mechanosensory-cued hatching (Jung and Warkentin, unpublished data). The constant ciliary circulation of the perivitelline fluid within *A. callidryas* eggs (Rogge and Warkentin, 2008; Warkentin et al., 2005) presumably stimulates the lateral line, and any change in this circulation pattern would therefore be perceptible to embryos.

The sensation of touch in adult frogs and tadpoles depends on cutaneous mechanoreceptors that are diverse and highly specialized (Catton, 1976; Fromy et al., 2008; Spray, 1976; Weston, 1970). A single mechanoreceptive afferent can encode more than one type of stimulus, for example temperature and texture (Hunt and McIntyre, 1960), as well as mechanical stimuli such as pressure and vibration (Ribot-Ciscar et al., 1989). Since all somatosensory neurons arise from precursor neural crest cells early in embryonic development, much prior to the development of the vestibular system (Jenkins and Lumpkin, 2017; Weston, 1970), pre-VOR *A. callidryas* embryos are likely to already have cutaneous mechanoreceptors. These could enable embryos to sense contact cues, through the membrane, as a probe or a predator touches the egg capsule. Moreover, if the inertia of embryos is higher than their surroundings, moving an egg might also change how strongly the embryos’ skin presses against the adjacent membrane, altering contact cues.

### Multiple mechanosensory cues in attacks and multiple mechanosensory systems

Several types of mechanosensory cues could occur in egg-predator attacks––and in egg-jiggling––including whole-egg motion, embryo motion within the capsule, and tactile contact that may deform egg-capsules or contact embryos through their perivitelline membranes (Fig. 9). Whole-egg motion occurs in vibration-playbacks, egg-jiggling, and predator attacks. This will activate the vestibular system as the embryo is passively accelerated along with its surrounding capsule. If the embryo remains in the same position relative to its capsule, whole-egg motion alone would likely not alter perivitelline fluid flow and seems unlikely to stimulate the lateral line or cutaneous touch receptors.

**Fig. 9.**
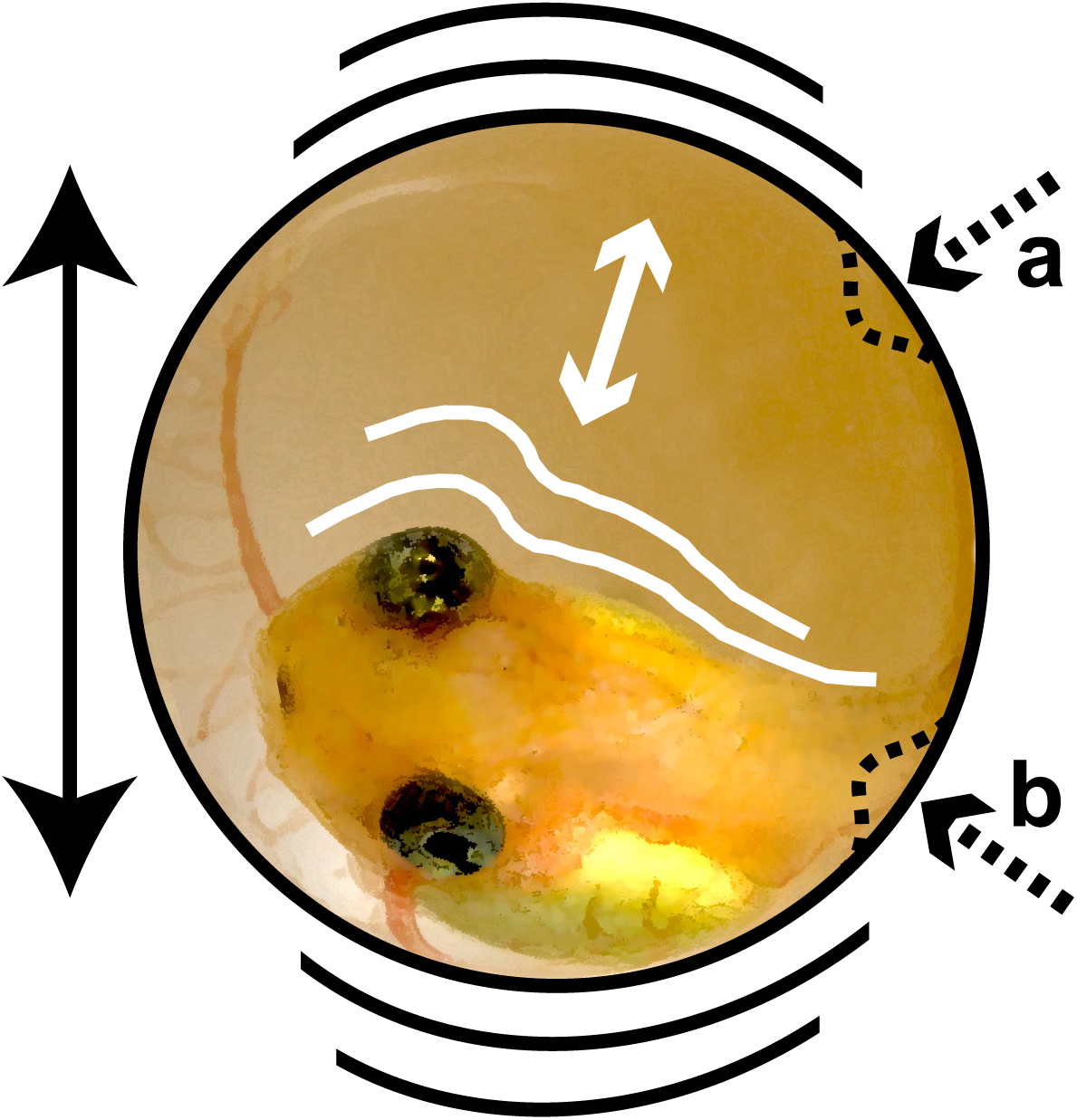
Types of mechanosensory cues in egg-predator attacks. Attacked embryos may experience whole-egg motion (solid black), embryo motion within the perivitelline chamber (white), and direct contact with eggs (dashed) that may **a.** deform egg capsules or **b.** touch embryos through their capsule.

Embryo motion within the capsule occurs when embryos are displaced in their perivitelline chamber as the capsule is moved. The inertia of the embryo likely differs from the surrounding fluid and capsule, such that the embryo may lag a bit behind as the egg accelerates around it. For instance, if the egg were accelerated up the embryo could be pressed against the bottom of the chamber, and if the egg were accelerated down the embryo could be lifted off the bottom. This could change both cutaneous stimulation and perivitelline fluid flow if the embryo’s body were sufficiently displaced within the capsule. Tactile contact occurs for a subset of eggs in predator attacks on, and tine-based vibration playbacks to, whole egg clutches, and for all eggs exposed to individual egg-jiggling stimulation. If the contact deforms egg capsules (e.g., dents or squashes them), even without contacting the embryo inside, it may change perivitelline fluid flow and lateral line input (Fig. 9). Contact with the embryos through the membrane would also directly stimulate cutaneous touch receptors.

Our egg-jiggling and vibration-playback experiments differed in several important ways. First, the jiggling stimulus represents a targeted attack on individual eggs rather than a generalized stimulus to whole clutches. Second, it was a more complex multimodal stimulus that combined whole egg motion with tactile elements and included both lateral and rolling movements. Since predators must touch eggs to eat them, risk of mortality in attacks is presumably higher for eggs receiving motion and contact cues than for those receiving motion cues alone. Both targeted jiggling of and predator attacks on individual eggs likely stimulate the vestibular system, the lateral line, and touch receptors in the skin. But other eggs in attacks and in vibration playbacks likely experience only whole-egg motion and vestibular stimulation. This variation in the cues available to embryos may contribute to the variation in individual responses and the different responses to vibration-playback and egg-jiggling stimuli. Moreover, in the jiggling series, embryos began showing VOR (thus, developing vestibular function) at earlier developmental stages compared to in the tine playback series (Fig. 6A vs. 5A), which may also have contributed to their earlier mechanosensory cued hatching.

### Ontogenetic changes in embryo use of multimodal mechanosensory cues

Whatever sensory system mediates hatching in egg jiggling for animals lacking VOR would presumably add to the stimulation experienced by older animals that have developed a functional vestibular system. Moreover, at a given stage of development, cues indicating greater risk should be more likely to elicit hatching. Thus, at the same stage, we expect an individually targeted “attack” stimulus to more strongly elicit hatching than a stimulus transmitted through the clutch. Consistent with this, stage 6 or 7 animals with strong VOR show a stronger hatching response to egg jiggling than to vibration playbacks to clutches. Nonetheless, if an animal has less-developed mechanoreceptors and cannot sense components of a stimulus, it will be limited in its risk-assessment ability. Stage 3 animals lacking VOR, and vestibular function, presumably receive just cutaneous and perhaps lateral line input in attacks and our mechanosensory stimuli. In contrast, stage 3–5 animals with low VOR likely also receive weak vestibular input; combining this with cutaneous and/or lateral line input may generate sufficient total stimulation to elicit hatching. Without additional input from another mechanosensory system, weak vestibular input may be insufficient to elicit hatching. Different types of mechanosensory cues likely stimulate different mechanoreceptor types, or combinations thereof, providing different and potentially synergistic or complementary information about risk. Thus, *A. callidryas* embryos may use multimodal mechanosensory cues to inform escape-hatching decisions, particularly at the onset of vibration-cued hatching when their mechanosensory systems are less developed.

We recently developed a new vibration playback system to generate whole-egg motion without tactile contact cues or egg-shape deformation, demonstrating that egg-motion alone is sufficient to induce hatching (Warkentin et al., 2019). A second playback-system component adds a tactile contact cue, which appears to synergize with motion to increase hatching of 4-day embryos (Fouilloux, Jung, Ospina, Snyder, & Warkentin unpublished). Lateral line blocking and/or vestibular system ablation experiments, in conjunction with vibration playbacks, would be useful to assess the individual and potentially interacting roles of these mechanosensory systems in the hatching decisions of *A. callidryas* embryos.

### Conclusion

Hatching is a developmentally critical behavior that immediately impacts survival in multiple ecological contexts. Environmentally cued hatching is widespread and well-documented in all three major clades of bilateria and, in many species, embryos respond to multiple different factors or contexts (Warkentin, 2011a). Physical disturbance of eggs is a particularly salient and common cue to hatch among embryos of fishes (Martin et al., 2011), amphibians (Buckley et al., 2005; Gomez-Mestre et al., 2008; Goyes Vallejos et al., 2018; Touchon et al., 2011; Warkentin, 1995; Warkentin, 2000; Warkentin, 2011b), and reptiles (Doody, 2011; Doody and Paull, 2013; Doody et al., 2012), as well as many invertebrates (Endo et al., 2018; Mukai et al., 2014; Oyarzun and Strathmann, 2011; Tanaka et al., 2016; Whittington and Kearn, 1988). Presumably, all the vertebrates that use physical disturbance as a hatching cue have vestibular systems and cutaneous mechanoreceptors, but only the fish and amphibians have lateral lines. Moreover, some of the contexts that induce hatching in vertebrates seem likely to provide only whole-egg motion cues. For instance, grunion embryos are tightly coiled within their eggs and pressed against the capsule wall at a stage when tumbling in waves elicits hatching (Martin et al., 2011; Speer-Blank and Martin, 2004). This embryo size and position seem likely to prevent passive displacement within the perivitelline chamber as eggs are moved. Pig-nosed turtle embryos hatch in response to a whole-egg motion stimulus presented via an electronic shaker in the laboratory (Doody et al., 2012). Neither the lateral line nor cutaneous sensing seem likely to play a role in these instances, suggesting the vestibular system could mediate motion-cued hatching responses in multiple–– perhaps many––vertebrate embryos. The mechanisms that enable, regulate, and inform hatching change developmentally, altering embryos’ capacities for behavioral responses to cues. Thus, information on embryos’ sensory development will clarify how and why development changes behavior. This research elucidates how changing sensory and behavioral abilities can affect an essential early behavior and reveals a fundamental mechanism underlying phenotypic plasticity at a critical life history switch point.

## Acknowledgments

This research was funded by the Smithsonian Tropical Research Institute, the National Science Foundation (IOS-1354072 to K.M.W. and J.G.M.) and Boston University, including grants from BU’s Undergraduate Research Opportunity Program to S.M.P.A. and S.J.K. It was conducted under permits SC/A-15-14 and SE/A-46-15 from the Panamanian Ministerio de Ambiente, STRI IACUC protocol 2014-0601-2017, and BU IACUC protocol 14-008. We thank Adrian Tanner for helping design and build the tadpole rotator, Nora Moscowitz and Angelly Vasquez for help with egg care in 2015, and Alina Chaiyasarikul and Adeline Almanzar for assistance measuring images to calculate VOR. We thank members of the Gamboa Frog Group at STRI and BU Egg Science Research Group for discussions of this research at many stages of its development.

## Author Contributions

J.G.M. and K.M.W. designed and built the tadpole rotator and J.G.M. provided vibrations engineering support for playbacks. S.M.P.A. and K.M.W. designed, and S.M.P.A. conducted, experiments Ia and Ib. J.J., S.J.K., and K.M.W. designed, and J.J. and S.J.K. conducted, experiments II and III. S.M.P.A, J.J. and S.J.K. measured VOR for all experiments. J.J. analyzed the data from all experiments. J.J. wrote the paper. All authors edited the paper.

